# RNA-sensing pattern-recognition receptors synergize with TLR2 or dectin-1 to trigger high production of IL-12p70 by human dendritic cells

**DOI:** 10.1101/2022.10.03.510527

**Authors:** Brian C. Gilmour, Alexandre Corthay, Inger Øynebråten

## Abstract

IL-12p70 is crucial for T helper 1 polarization and the generation of type 1 immunity that is required to fight cancer and intracellular pathogens. Therefore, strategies to optimize the production of IL-12p70 by dendritic cells (DCs) may significantly improve the efficacy of vaccines and immunotherapies for cancer. However, the rules governing the production of IL-12p70 remain obscure. Here, we stimulated pattern recognition receptors (PRRs) representing all five families of PRRs, to evaluate their ability to elicit high production of IL-12p70 by human DCs. We used ten well-characterized agonists and stimulated human monocyte-derived DCs *in vitro* with either single agonists or 26 different combinations. We found that poly(I:C), which engages the RNA-sensing PRRs TLR3 and/or MDA5, was the only agonist that could elicit IL-12p70 production when used alone. Combinations of agonists of cell surface and intracellular PRRs were found to synergize to induce high IL-12p70 production, given that the combination included poly(I:C) or resiquimod, which both are agonists of intracellular, RNA sensing PRRs (TLR3/MDA5 and TLR7/8, respectively). Our data show that production of high IL-12p70 is strictly controlled, which is different from what we observed for IFNβ, whose production could be elicited by several intracellular PRRs. In conclusion, we identified six different combinations of PRR ligands able to induce high IL-12p70 production by human DCs. The identified synergistic PRR ligand combinations may represent strong adjuvant candidates in particular for therapeutic cancer vaccines.

## Introduction

In the 3-signal model of naïve T cell activation, T cells are activated upon recognition of cognate antigen wherein antigen-presenting cells (APCs) provide signal 1 and 2 by presenting the peptide-MHC complex and co-stimulatory molecules, respectively, to T cells^1-5^. Signal 3 consists of cytokines, and IL-12p70 is crucial for the polarization of naïve CD4^+^ T cells into T helper 1 (Th1) cells^6-8^. A successful immune response against cancer is typically associated with a strong Th1 and CD8^+^ T cell response, *i.e*. type 1 immunity^9-11^. In vaccines consisting of proteins or peptides from pathogens or neoantigens in cancer, molecular patterns which normally would evoke a pro-inflammatory response are lacking, and there is a need for adjuvants that can elicit cytokines important for strong type 1 immunity. Strategies that can optimize the production of IL-12p70 by dendritic cells (DCs) may significantly improve the efficacy of vaccines and immunotherapies for cancer.

Pattern recognition receptors (PRRs) can contribute to all three signals of the 3-signal model since APCs stimulated with PRR agonists may up-regulate antigen-presentation and increase the expression of co-stimulatory molecules as well as signal 3 cytokines. Thus, there is a great interest in PRRs as targets of adjuvants in vaccines^12^. However, with the exception of toll-like receptor 3 (TLR3), the ability of different PRRs to trigger IL-12p70 production remains poorly characterized^12^. Several transcription factors are likely involved in IL-12p70 production of which interferon regulatory factors (IRFs) and NF-κB are the most studied^13-17^. IL-12p70 is a heterodimer of two disulfide-linked protein units, p35 and p40, which are recognized by the β2 and β1 chain, respectively, of the IL-12 receptor^18,19^. The p35 unit is considered the limiting factor in IL-12p70 production, but how the p35 gene is activated and thereby IL-12p70 production is regulated, remains obscure^18,20,21^. IL-12p40 can be present as a monomer, homodimer, or a heterodimer where the p35 chain may be substituted with p19 to form IL-23^22^. Therefore, quantitative data on the p40 unit/IL-12p40, which is presented in numerous studies, is not a reliable measure of IL-12p70. Only specific measurement of the IL-12p70 heterodimer is representative of the Th1 polarizing cytokine IL-12p70.

PRRs are divided into five families based on protein domain homology: *i*) TLRs; *ii*) absent in melanoma 2 (AIM2)-like receptors (ALRs); *iii*) C-type lectin receptors (CLRs); *iv*) nucleotide-binding oligomerization domain (NOD)-like receptors (NLRs); and *v*) retinoic acid-inducible gene (RIG)-I-like receptors (RLRs). The TLRs constitute the best characterised family of PRRs and recognize a variety of ligands including lipopolysaccharide (LPS), lipoprotein, double stranded (ds) and single stranded (ss) RNA, as well as unmethylated CpG-containing dsDNA^23,24^. CLRs bind carbohydrate moieties and sense various pathogens including fungi, as well as self-ligands such as dying cells^25-29^. ALRs, NLRs, and RLRs are all localized in the cytosol, but are engaged by different ligands: NLRs recognize peptidoglycans whereas ALRs and RLRs recognize DNA and RNA, respectively^30^.

Until now, only two PRR agonists, monophosphoryl lipid A and CpG-B 1018 which stimulate TLR4 and TLR9, respectively, have been included in licensed vaccines towards infectious diseases^31-34^. In cancer immunotherapy, combined stimulation of TLR3 and TLR7/8 has been used in clinical studies to activate monocyte-derived DCs (moDCs)^35,36^. Finally, TLR3 agonists have been tested in numerous preclinical studies and some have been transferred to clinical studies^37^. However, the potential of other PRR family members is poorly characterized.

In this study, we used established PRR agonists to engage receptors among all five families of PRRs. The ligands were added to cultures of moDCs, and the signal 3 cytokine IL-12p70, the antiviral cytokine IFNβ, and the immunosuppressive cytokine IL-10 were quantified in the cell culture media. DC activation was also assessed by measurement of cell surface expression of CCR7, HLA-DR, and the co-stimulatory molecules CD80 and CD86. In total, we used ten ligands to stimulate 11 different PRRs. When DCs were treated with single agonists, only poly(I:C), a double stranded RNA molecule that binds to TLR3 and MDA5, induced notable IL-12p70 production, whereas resiquimod, a ligand of TLR7/8 induced IL-12p70 in some experiments. Poly(I:C) and R848 in combination with ligands of the cell surface receptors dectin-1 and TLR2:1, respectively, synergized and induced high levels of IL-12p70. The tested PRR ligands could be divided into three groups depending on whether they triggered production of both IL-12p70 and IFNβ, only one of the two or neither. Our data suggest that high production of IL-12p70 is strictly controlled.

## Methods

### Positive selection of human monocytes

Monocytes were isolated from buffy coats obtained from the blood bank at Oslo University Hospital, Oslo, Norway, and approved for use by the Norwegian Regional Committee for Medical and Health Research Ethics, REK no. 2019/113. The buffy coats were diluted 1:1 with sterile phosphate buffered saline without Mg^2+^ and Ca^2+^ (PBS^-/-^) containing 2 % fetal bovine serum (FBS) (Biowest, #S181BH), before gently being added on top of Lymphoprep^™^ (Progen, #1114545) in 50 mL tubes in volumes recommended by the provider. The tubes were centrifuged at 800 g for 20 min at room temperature (RT) with the brake disabled. Peripheral blood mononuclear cells (PBMCs) located at the interface of the plasma and Lymphoprep layers were collected and washed twice in PBS^-/-^ with 2 % FBS by centrifugation (400 g, 7 min at RT) to remove remnants of Lymphoprep. The pelleted PBMCs were re-suspended in PBS^-/-^ with 2 % FBS and passed through a 30 μm filter (Miltenyi, #130-041-407) to remove cell clumps and debris. Next, the monocytes were positively selected from the PBMCs by magnetic-activated cell sorting (MACS) technology using CD14^+^ MicroBeads (Miltenyi, #130-050-201) according to the manufacturer’s instructions: PBMCs, per 10^7^ cells, were re-suspended in 80 μL cold MACS buffer (PBS^-/-^ with 10 % FBS and 1 % EDTA) and mixed with 20 μL MicroBeads, to a maximum volume of 2 mL in the presence of higher cell numbers. The mixture of PBMCs and MicroBeads was vortexed before incubation for 15 min at 4 °C. Next, after washing of the cells, maximum 1 × 10^8^ labeled cells were applied onto an LS column (Miltenyi, #130-042-401) placed in a MACS magnet separator (Miltenyi, #130-042-109). Unlabeled cells were washed out before the magnetically labelled cells were harvested in 5 mL PBS^-/-^ with 2 % FBS by use of a plunger. Staining with an APC/Cy7-conjugated anti-human CD14 antibody (clone HCD14) followed by flow cytometry, showed that >95% of the positively selected cells were monocytes (n=3). The positively selected monocytes were either frozen in FBS containing 10 % DMSO (PanReac AppliChem, #67-68-5) or immediately differentiated by GM-CSF and IL-4 as described below.

### Differentiation protocol for human monocyte-derived dendritic cells

On day 0, 2-2.5 × 10^6^ frozen or freshly isolated monocytes were seeded in 10 cm non-tissue culture treated dishes (VWR, #734-2796) in 10 mL RPMI 1640 (Biowest, #L0500) with 10 % FBS (Biowest, #S181BH), 1 % Penicillin-Streptomycin (Biowest, #L0022), 100 ng/mL GM-CSF (PeproTech, #300-03) and 20 ng/mL IL-4 (PeproTech, #200-04). The cells were cultivated at 37 °C and 5 % CO_2_. On day 3, medium was replenished by adding 10 mL fresh medium containing 100 ng/mL GM-CSF and 20 ng/mL IL-4. At day 5, the cells were harvested and seeded out for use in experiments. Non-adhesive cells were collected directly from the medium. Loosely adherent cells were harvested by applying 5 mL cold PBS^-/-^ to the dish for 5-10 min at RT before harvesting the cells by pipetting the PBS^-/-^ carefully up and down on the plate 3-5 times.

### Stimulation with pattern recognition receptor ligands

Harvested day 5 moDCs were plated in 96-well plates (Corning, #3595) (120 000 cells in 100 μL/well) for quantification of released cytokines, and in non-tissue culture treated 6-well plates (Corning, #3736) (500 000 cells in 1.4 mL/well) for flow cytometry analysis of cell surface molecules. RPMI 1640 with 10 % FBS, 1 % Penicillin-Streptomycin, 100 ng/mL GM-CSF, and 20 ng/mL IL-4 was used as medium. After cultivation for 24 h at 37 °C and 5 % CO_2_, PRR ligands (Table 1) were added to the moDC cultures (in 6x concentration to the 96-well plates to a total volume of 120 μL). To target cytosolic PRRs, LyoVec^™^ reagent (Invivogen, #lyec-12) was used to transfect the ligands 3p-hpRNA (RIG-I), poly(I:C) (MDA5/TLR3), and p(dA:dT) (cGAS/ALR) into the cytosol according to manufacturer’s protocol. Briefly, the desired ligand was diluted in LyoVec^™^ and incubated for at least 15 min to a maximum of 1 h. Next, the mixture was combined 1:1 with RPMI 1640 containing 10 % FBS, 1 % Penicillin-Streptomycin and, alternatively another PRR ligand. MoDCs were then incubated with the mixture for 24 h.

**Table 1.**
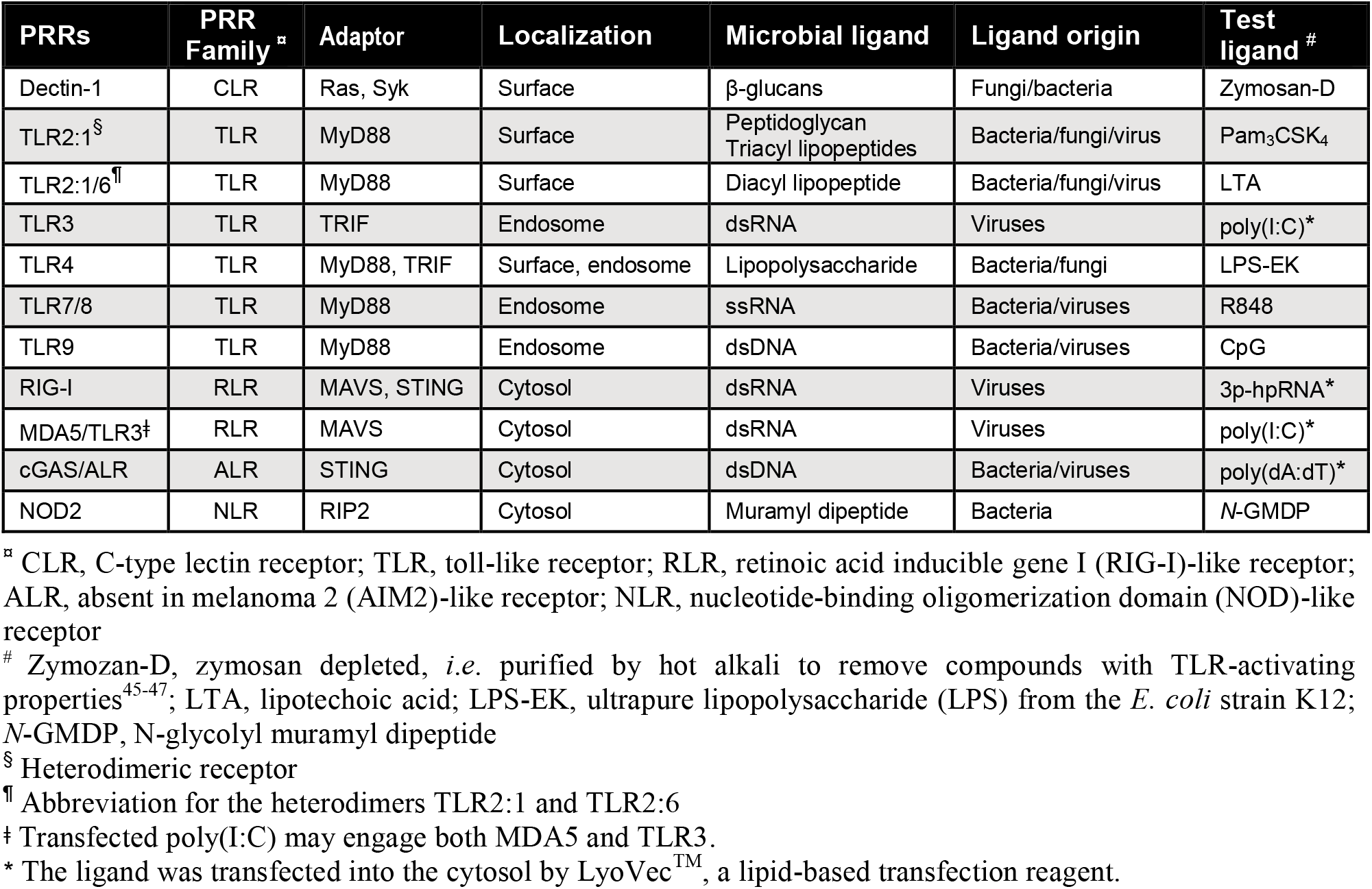
Pattern recognition receptors (PRRs) and the respective agonists selected for this study.

| PRRs | PRR Family <sup>□</sup> | Adaptor | Localization | Microbial ligand | Ligand origin | Test ligand <sup>#</sup> |
| --- | --- | --- | --- | --- | --- | --- |
| Dectin-1 | CLR | Ras, Syk | Surface | β-glucans | Fungi/bacteria | Zymosan-D |
| TLR2:1<sup>§</sup> | TLR | MyD88 | Surface | Peptidoglycan<br>Triacyl lipopeptides | Bacteria/fungi/virus | Pam<sub>3</sub>CSK<sub>4</sub> |
| TLR2:1/6<sup>¶</sup> | TLR | MyD88 | Surface | Diacyl lipopeptide | Bacteria/fungi/virus | LTA |
| TLR3 | TLR | TRIF | Endosome | dsRNA | Viruses | poly(I:C)* |
| TLR4 | TLR | MyD88, TRIF | Surface, endosome | Lipopolysaccharide | Bacteria/fungi | LPS-EK |
| TLR7/8 | TLR | MyD88 | Endosome | ssRNA | Bacteria/viruses | R848 |
| TLR9 | TLR | MyD88 | Endosome | dsDNA | Bacteria/viruses | CpG |
| RIG-I | RLR | MAVS, STING | Cytosol | dsRNA | Viruses | 3p-hpRNA* |
| MDA5/TLR3<sup>‡</sup> | RLR | MAVS | Cytosol | dsRNA | Viruses | poly(I:C)* |
| cGAS/ALR | ALR | STING | Cytosol | dsDNA | Bacteria/viruses | poly(dA:dT)* |
| NOD2 | NLR | RIP2 | Cytosol | Muramyl dipeptide | Bacteria | N-GMDP |

<sup>□</sup> CLR, C-type lectin receptor; TLR, toll-like receptor; RLR, retinoic acid inducible gene I (RIG-I)-like receptor; ALR, absent in melanoma 2 (AIM2)-like receptor; NLR, nucleotide-binding oligomerization domain (NOD)-like receptor
<sup>#</sup> Zymosan-D, zymosan depleted, *i.e.* purified by hot alkali to remove compounds with TLR-activating properties<sup>45-47</sup>; LTA, lipotechoic acid; LPS-EK, ultrapure lipopolysaccharide (LPS) from the *E. coli* strain K12; N-GMDP, N-glycolyl muramyl dipeptide
<sup>§</sup> Heterodimeric receptor
<sup>¶</sup> Abbreviation for the heterodimers TLR2:1 and TLR2:6
<sup>‡</sup> Transfected poly(I:C) may engage both MDA5 and TLR3.
\* The ligand was transfected into the cytosol by LyoVec<sup>TM</sup>, a lipid-based transfection reagent.

For cytokine quantification, the culture medium was harvested from 96-well plates. Cells and debris were removed from the culture medium by centrifugation two times at 4 °C, first at 400 g for 7 min, and then at 1 000 g for 15 min. The supernatants were then frozen and stored at -80 °C until analysis.

### ELISA

IFNβ, IL-10, and IL-12p70 were quantified by use of ELISA kits (Supplementary Table 2), carried out in half-area, flat-bottom 96-well plates (Corning, #3690). Excluding PBS^-/-^, all necessary diluents and buffers were from the DuoSet^®^ Ancillary Reagent Kit 2 (R&D Systems, #DY008). The assays were performed at RT unless otherwise noted, and all steps were followed by washing using 200 μL/well of wash buffer (R&D Systems) three times. The desired capture antibody (Supplementary Table 2) was diluted to the concentration recommended by the provider, and 50 μL of the antibody was applied per well before incubation overnight. Next, the wells were blocked using 150 μL/well of reagent diluent (PBS^-/-^ with 1 % bovine serum albumin (BSA)) for at least 1 h. Afterwards, 50 μL/well of samples, or standard in 2-fold dilution series were added in duplicates. The plate was incubated for 2 h and washed, before incubation for another 2 h with 50 μL/well of biotinylated detection antibody. Streptavidin-HRP diluted 1:40 (50 μL/well) was incubated for 20 min. After washing, 50 μL/well of a 1:1 solution of H_2_O_2_ (R&D Systems, part #895000) and tetramethylbenzidine (R&D Systems, part #895001) was added for approx. 20 min before the reaction was stopped by addition of 25 μL/well of 2N H_2_SO_4_. Absorbance was measured at 450 nm and 540 nm (reference wavelength) using an Epoch^™^ Microplate Spectrophotometer, with the 540 nm readings subtracted from the 450 before analysis. Standard curves were made by fitting a four parameter logistic (4PL) curve model to the data.

### Flow cytometry of surface molecules

Harvested cells were kept on ice for approx. 15 min, before 200 000 cells re-suspended in flow buffer (PBS^-/-^ with 10 % FBS) were portioned out per staining condition. Fc-receptors were blocked by adding 2.5 μL Human TruStain^™^ FcX (BioLegend, #101320) per 50 μL of 200 000 cells, and after incubation for 30 min on ice, the cells were pelleted by centrifugation (400 g, 7 min at 4 °C). Next, the moDCs were re-suspended in 50 μL flow buffer containing 2.5 μL of fluorophore-conjugated antibody (Supplementary Table 3). Irrelevantly targeted antibodies, but otherwise matched to the specific antibodies with regard to fluorophore, concentration, isotype, and species were used as isotype controls (Supplementary Table 4). Following incubation for 20 min in the dark on ice, unbound antibody was removed by washing in flow buffer. Finally, the moDCs were re-suspended in 200 μL flow buffer. Immediately before analysis on a BD LSRFortessa^™^ flow cytometer, 2.5 μL propidium iodide (PI) (BioLegend, #421301) was added enabling the elimination of dead cells. eBioscience^™^ UltraComp eBeads (Invitrogen, #01-2222-42) were used to adjust the settings for compensation. Isotype controls and unstained cells were used to set the threshold for positive staining. Data was analyzed using FlowJo V10 software (Becton Dickinson and Company), and geometric mean fluorescent intensity (GMFI) values were exported for further analysis.

## Results

### Only a few PRR agonists trigger production of IL-12p70 and IFNβ by human monocyte-derived DCs when used as single ligands

To investigate the effect of different PRR agonists on the production of IL-12p70 and IFNβ by human DCs, we isolated CD14^+^ monocytes from healthy blood donors and generated moDCs by culturing the cells *in vitro* for 6 days with GM-CSF and IL-4 according to an established protocol^35,38^. Flow cytometry analysis showed that CD14 expression was down-regulated during differentiation to moDCs, which acquired a typical CD14^lo^, CD11c^hi^, and HLA-DR^hi^ DC phenotype (Supplementary Fig. 1, Supplementary Fig. 2). Ten PRR agonists were selected for this study and include ligands engaging PRRs of all five PRR families (see Table 1 with additional information provided in Supplementary Table 1). The moDCs were stimulated *in vitro* with the PRR agonists for 24 h, and the cytokines IL-12p70, IFNβ and IL-10 released into the cell medium were quantified by ELISA.

In the first experimental setting, moDCs were stimulated with ten single PRR ligands, and unstimulated moDCs were used as control (Fig. 1). Barely detectable amounts of IL-12p70 were released from unstimulated moDCs (Fig. 1a). Strikingly, the TLR3 and MDA5 agonist poly(I:C) was the only ligand able to induce notable and consistent IL-12p70 production by moDCs (Fig. 1a, Supplementary Table 5). In addition, the TLR7/8 agonist R848 was found to induce low levels of IL-12p70 in some, but not all, experiments. Furthermore, three single PRR agonists were able to induce secretion of IFNβ by moDCs: the TLR3 and MDA5 agonist poly(I:C), the RIG-1 agonist 3p-hpRNA, and the cGAS agonist poly(dA:dT) (Fig. 1b, Supplementary Table 6). In contrast to what we observed for IL-12p70 in some experiments, R848 did not induce detectable IFNβ production (Fig. 1b). Taken together, the data show that secretion by human moDCs of IL-12p70 and IFNβ is only triggered by some PRRs. Moreover, there is only a partial overlap between the PRRs that induced notable production of IL-12p70 (TLR3, MDA5) and IFNβ (TLR3, MDA5, RIG-1, cGAS), suggesting that there are different rules for production of IL-12p70 and IFNβ.

**Figure 1.**
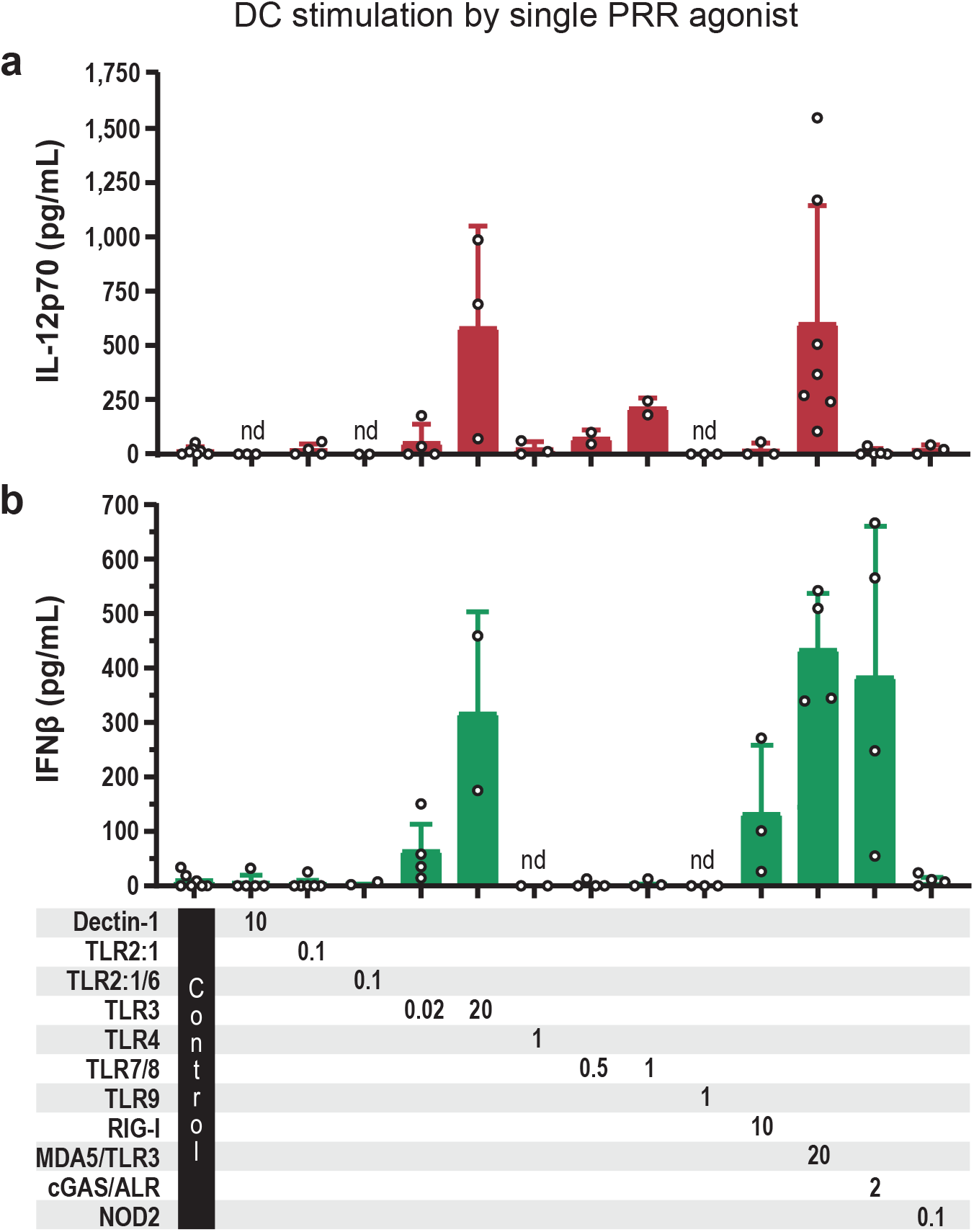
IL-12p70 and IFNβ released from moDCs stimulated with single ligands engaging the indicated PRRs. Cultures of moDCs were stimulated for 24 h with single PRR-ligands before **a** IL-12p70, and **b** IFNβ were measured in the cell culture supernatants. Unstimulated moDCs were used as control. Each circle shows the cytokine concentration from one experiment, and the bars present the mean with SD (n=2-7). The numbers below the bars indicate the ligand concentrations in μg/mL. ALR, absent in melanoma 2 (AIM2)-like receptor; CLR, C-type lectin receptor; NLR, nucleotide-binding oligomerization domain (NOD)-like receptor; RLR, retinoic acid-inducible gene (RIG)-I-like receptor; TLR, toll-like receptor. nd, not detectable.

### Six different combinations of PRR agonists, either containing poly(I:C) or R848 engaging RNA-sensing PRRs, elicit high levels of IL-12p70

A pathogen is likely recognized by several PRRs, and we therefore stimulated the cell surface PRRs dectin-1 or TLR2 in combination with an intracellular PRR (Fig. 2a, b). In other experiments, combinations of intracellular PRRs were stimulated (Fig. 2c, d). When dectin-1 was stimulated by use of zymosan-D, only the combinations containing ligands of TLR3, MDA5, or TLR7/8 resulted in moderate-to-high amounts of IL-12p70 (Fig. 2a). Based on the levels of IL-12p70 that were generated, we considered dectin-1+TLR7/8 to be the most promising combination and named it C1. Whereas dectin-1 and TLR7/8 separately induced non-detectable and at the most, low levels of IL-12p70, respectively, the combination C1 resulted in >2000 pg/mL IL-12p70, revealing a synergistic effect upon stimulation of the two PRRs (Fig. 1a, 2a).

**Figure 2.**
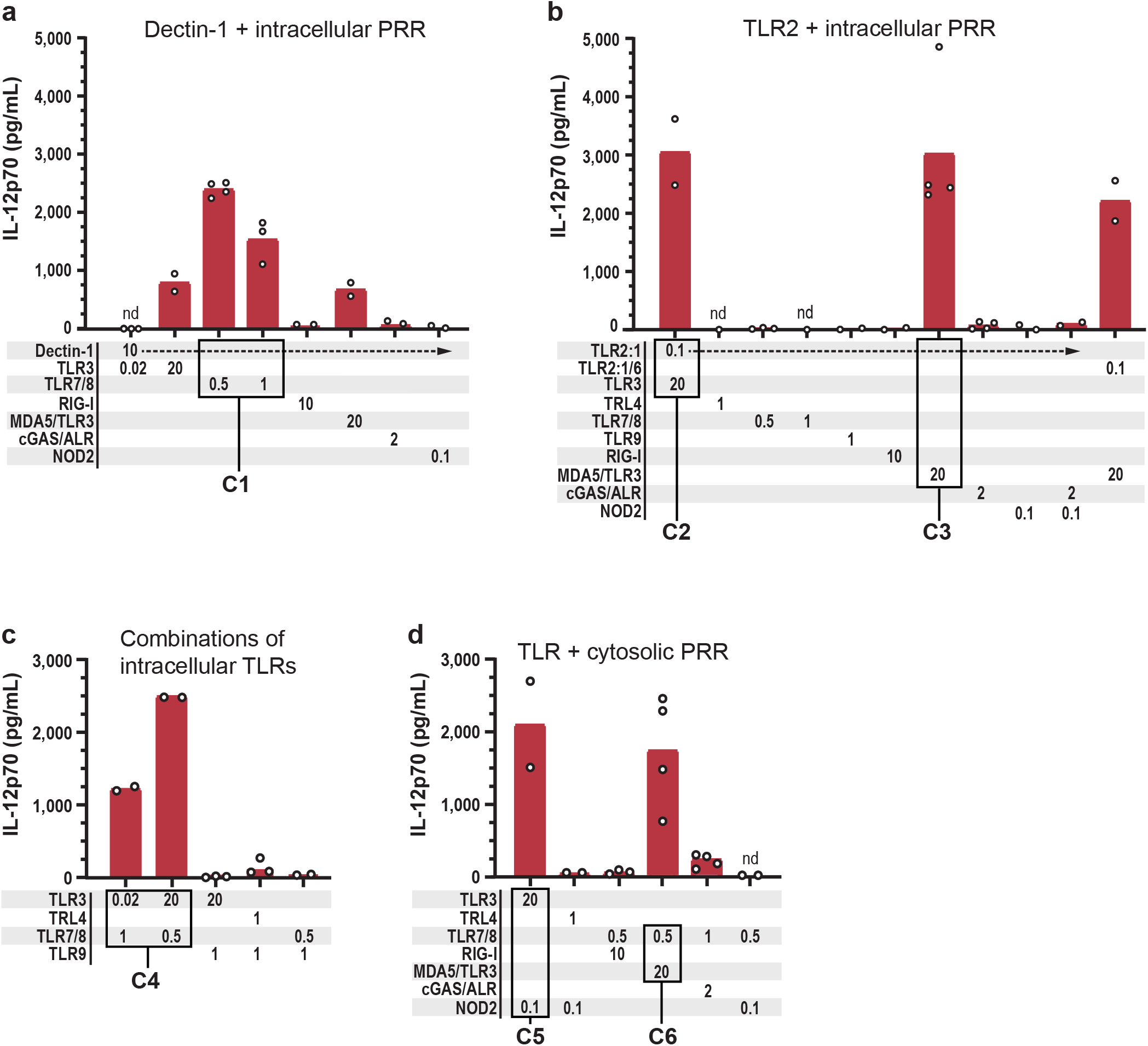
The TLR3 and MDA5 agonist poly(I:C) can be combined with ligands of cell surface PRRs as well as intracellular PRRs to elicit high IL-12p70 production. IL-12p70 was quantified in the cell culture media of moDCs stimulated for 24 h with ligands of the indicated PRRs. Various combinations of PRRs were stimulated: The cell surface PRRs **a** dectin-1 or **b** TLR2 in combinations with intra-cellular PRRs; **c** combinations of intracellular TLRs; and **d** intracellular TLR + cytosolic PRR. The ligand concentrations in μg/mL are given below the bars. Each circle presents the concentration from one inde-pendent experiment, and bars represent the mean (n=2-4). C1-C6 highlight the PRR combinations which resulted in high IL-12p70 levels. nd, not detectable.

Pam_3_CSK_4_-mediated stimulation of the cell surface PRR TLR2:1 induced high production of IL-12p70 only when it was combined with poly(I:C), either non-transfected (TLR3) or transfected (MDA5/TLR3) (Fig. 2b). The combinations are designated C2 and C3. Stimulation of TLR2:1 in combination with the remaining PRRs, including TLR7/8, resulted in barely detectable or non-detectable levels of IL-12p70 (Fig. 2b). Thus, whereas dectin-1 could be stimulated together with TLR3, MDA5, or TLR7/8 to trigger production of IL-12p70, TLR2:1-stimulation resulted in high levels of IL-12p70 only when it was combined with poly(I:C) (TLR3, MDA5).

Combined stimulation of intracellular PRRs by use of poly(I:C) (TLR3) + R848 (TLR7/8) induced high production of IL-12p70 (combination C4; Fig. 2c). We tested two different doses of the ligands and found that 20 μg/mL and 0.5 μg/mL of poly(I:C) and R848, respectively, induced 2-fold more IL-12p70 than 0.02 μg/mL poly(I:C) + 1 μg/mL R848. TLR3 stimulation could also be combined with NOD2 (C5), and TLR7/8 stimulation could be combined with MDA5/TLR3 (C6) to induce high IL-12p70 production (Fig. 2d). Taken together, poly(I:C), the ligand of TLR3 and MDA5, could be combined with any of tested ligands of intracellular PRRs (with the exception of TLR9) and elicit production of IL-12p70.

### The combinations C1 (dectin-1+TLR7/8), C3 (TLR2:1+MDA5/TLR3), and C4 (TLR3+TLR7/8) synergized to induce high production of IL-12p70

We performed side-by-side comparisons to identify whether combined stimulation of PRRs in comparison with stimulation of separate PRRs, significantly increased the level of IL-12p70. We tested three of the combinations which in the first experiments demonstrated high IL-12p70 production (>1500 pg/mL): C1 (dectin-1+TLR7/8), C3 (TLR2:1+MDA5), and C4 (TLR3+TLR7/8) (Fig. 2a-c). Stimulation of TLR4+IFNγ receptor as well as the low poly(I:C) concentration of C4, which has been in clinical use^36^, were included for comparison (Fig. 3). All the tested combinations C1, C3, and C4 showed a synergistic effect on the level of IL-12p70 (Fig. 3). Synergy was most pronounced for dectin-1+TLR7/8 (C1), which in comparison to the separate PRRs increased the IL-12p70 level 25-fold and 47-fold, respectively. TLR2:1+MDA5/TLR3 (C3) induced more IL-12p70 than TLR3+TLR7/8 (C4), and all three combinations elicited >2-fold more IL-12p70 than TLR4+IFNγR (Fig. 3). In summary, selected PRRs could be combined to synergistically increase the IL-12p70 level. Thus, given the presumption that high levels of IL-12p70 are beneficial for development of Th1 and CD8^+^ T cell responses, the identified combinations of two PRR agonists have greater potential than one ligand as vaccine adjuvant.

**Figure 3.**
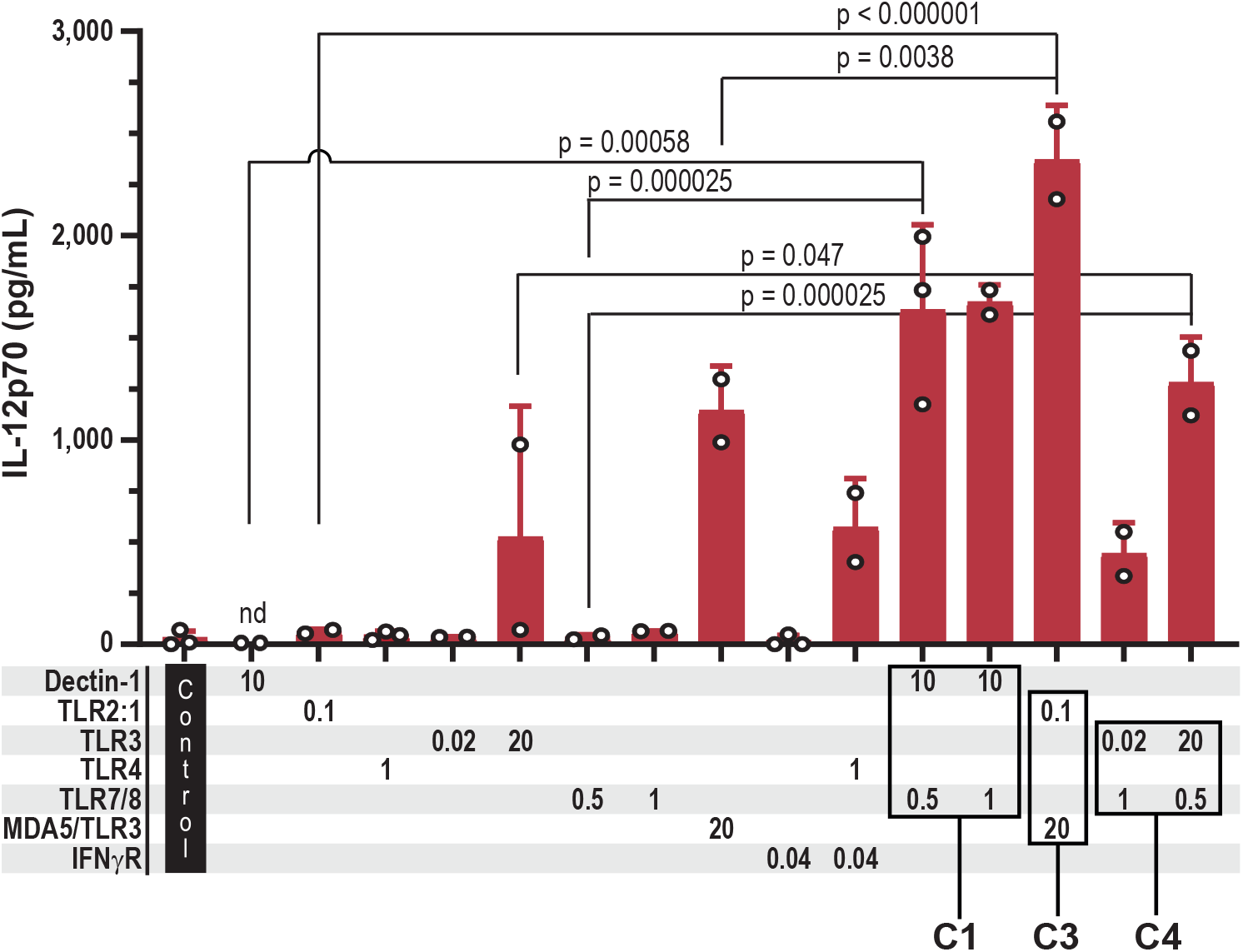
Combined stimulation with two PRR ligands synergistically increased IL-12p70 production. Cultures of moDCs were stimulated for 24 h with the previously identified ligand combinations (C1, C3, and C4) along with the respective single ligands. Unstimulated moDCs were set up as control. IL-12p70 was quantified in the cell culture supernatants, and each circle shows the concentration from one experiment. The ligand concentrations in μg/mL are indicated below the bars. Bars present the mean with SD (n=2-3). P values < 0.05 were considered statistically significant, and were calculated by ANOVA using the Šidák test to correct for multiple comparisons. IFNγ was used to activate the IFNγ receptor (IFNγR). nd, not detectable.

### Several combinations of PRR agonists induce IFNβ production

Next, we quantified IFNβ released in response to the same combinations that were tested for the ability to induce IL-12p70 production (Fig. 4). The combination C1 (dectin-1+TLR7/8), which triggered high production of IL-12p70 (Fig. 2a, 3), resulted in minor to non-detectable levels of IFNβ (Fig. 4a). However, stimulation of dectin-1 could be combined with stimulation of several other intracellular PRRs - TLR3, RIG-I, MDA5, or cGAS - to elicit IFNβ production (Fig. 4a, b). Moreover, both C2 (TLR2:1+TLR3) and C3 (TLR2:1+MDA5/TLR3), as well as stimulation of TLR2:1 in combination with RIG-1 or cGAS induced IFNβ production (Fig. 4b). Thus, IFNβ was produced when stimulation of the cell surface PRRs dectin-1 or TLR2:1 was combined with stimulation of an intracellular PRR that alone was able to elicit IFNβ production (Fig. 1b, 4a, b).

**Figure 4.**
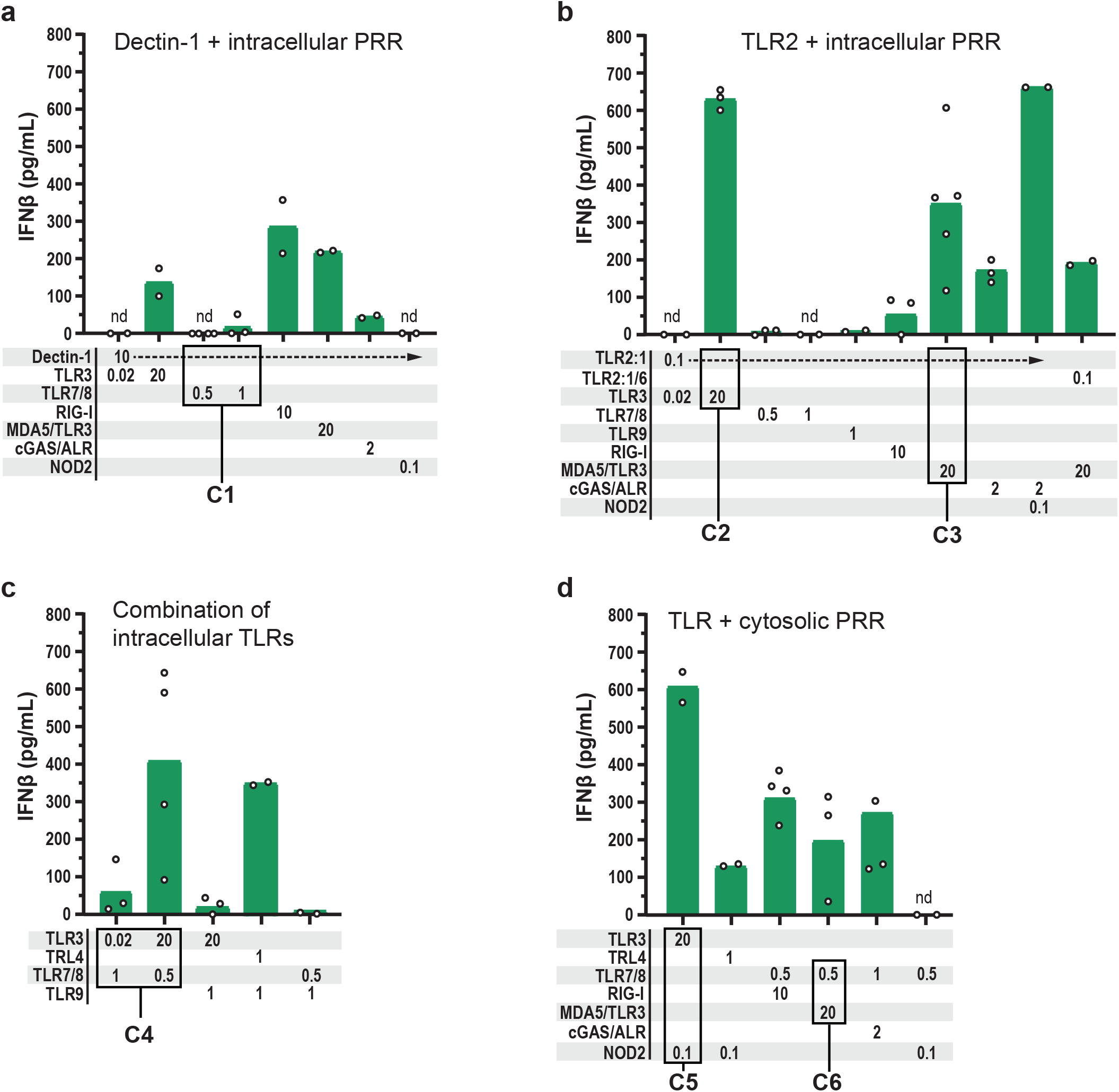
Several combinations of PRR ligands induced substantial levels of IFNβ. The indicated PRRs were stimulated for 24 h before IFNβ was quantified in the cell culture media of moDCs. Various combinations of PRRs were stimulated: The cell surface PRRs **a** dectin-1 or **b** TLR2 in combination with intracellular PRRs; **c** combinations of intracellular TLRs; and **d** intracellular TLR + cytosolic PRR. The concentrations of the PRR ligands are indicated in μg/mL below the bars. Each circle represents the IFNβ concentration from one experiment, and bars show the mean (n=2-5). The combinations C1-C6 induced high production of IL-12p70. nd, not detectable.

IFNβ was also produced in response to several of the combinations which stimulated intracellular PRRs, such as C4 (TLR3+TLR7/8), C5 (TLR3+NOD2), and C6 (TLR7/8+MDA5/TLR3) (Fig. 4c, d). Taken together, the data presented in Fig. 4 show that the combinations which included an intracellular PRR sensing DNA or RNA elicited IFNβ production. Among the combinations C1-6, which induced high levels of IL-12p70, all except C1 induced substantial production of IFNβ.

### Stimulation with a single PRR ligand is sufficient to induce high production of IFNβ

Finally, we tested whether the combinations dectin-1+TLR7/8 (C1), TLR2:1+MDA5/TLR3 (C3), or TLR3+TLR7/8 (C4) could act in synergy and increase the level of IFNβ (Fig. 5). In addition, TLR4 was stimulated in combination with the IFNγ-receptor, but the combination gave barely detectable levels of IFNβ (Fig. 5). In accordance with what we had observed already (Fig. 4a), stimulation of dectin-1+TLR7/8 (C1) resulted in IFNβ levels similar to that measured in media from unstimulated moDCs (Control), *i.e*. the levels were minor (Fig. 5). Stimulation of TLR2:1+MDA5/TLR3 (C3) or TLR3+TLR7/8 (C4) generated IFNβ levels in the same range as stimulation of MDA5 or TLR3 with poly(I:C) (Fig. 5). Thus, the combinations C1, C3, and C4 showed no synergistic effect on the level of IFNβ.

**Figure 5.**
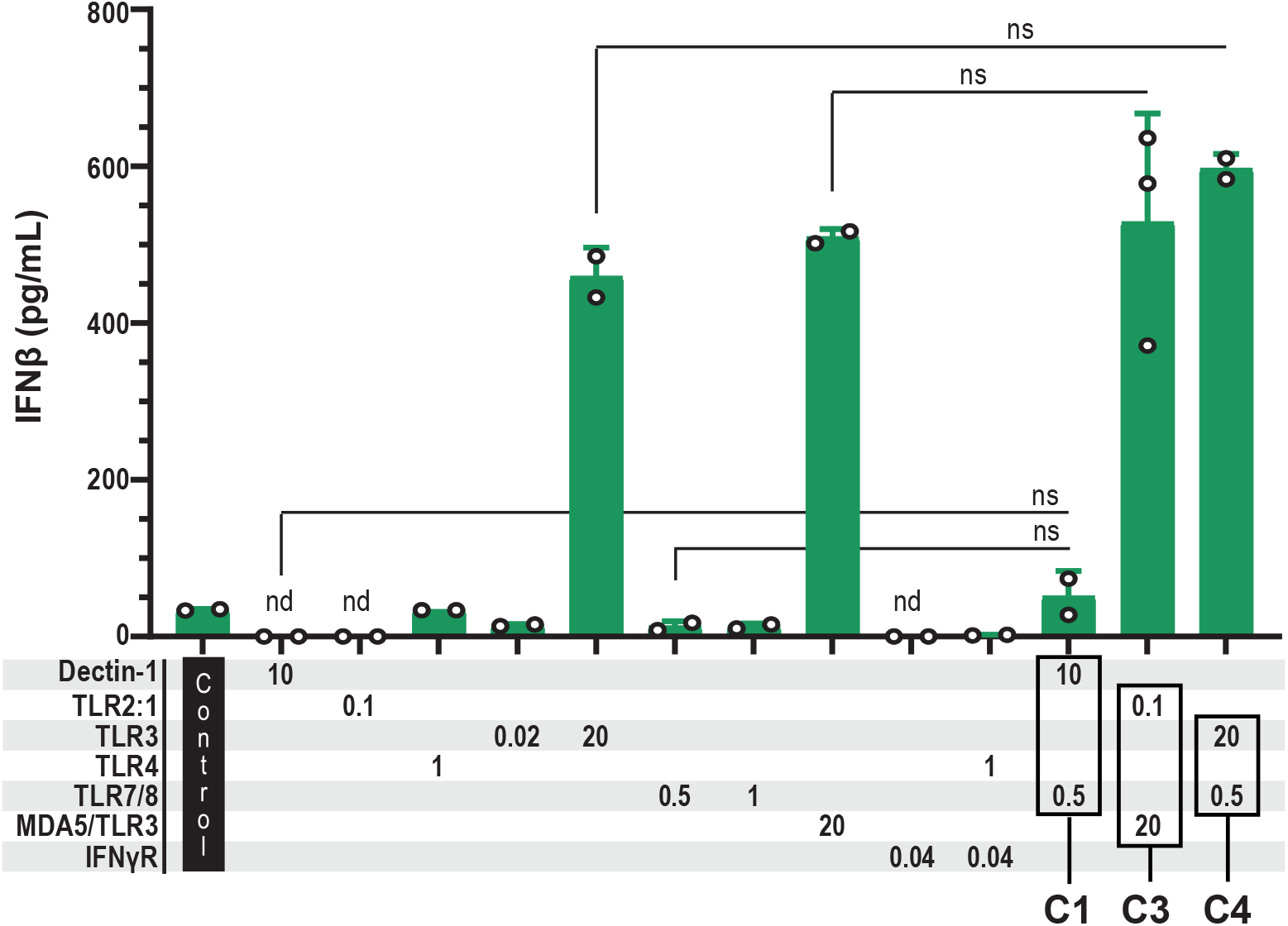
Stimulation with two PRR ligands elicited IFNβ levels similar to that of the respective single ligands. Cultures of moDCs were stimulated for 24 h with ligand combinations C1, C3, and C4 along with the respective single ligands, before IFNβ was quantitated in the cell culture supernatants. The used PRR ligand concentrations are indicated in μg/mL below the respective bars. Each circle shows the IFNβ concentration of one experiment, and bars represent the mean with SD (n=2-3). P values were calculated by ANOVA using the Šidák test to correct for multiple comparisons, and p values below 0.05 were considered statistically significant. Control, unstimulated moDCs. nd, not detectable; ns, not significant.

### Most PRR agonists reduced the level of secreted IL-10

Because IL-10 is an anti-inflammatory cytokine that can inhibit T cell activation and memory development, we examined whether a selection of PRRs upon stimulation altered the level of IL-10 produced by moDCs (Fig. 6). The highest levels of IL-10 were measured in cell culture media from unstimulated moDCs and from moDCs stimulated with the TLR2:1 ligand. In contrast, the remaining PRR ligands, either alone or in combinations, resulted in IL-10 levels which were 2.6 to 25-fold lower than in unstimulated moDCs (Fig. 6). In summary, apart from TLR2:1, stimulation of the selected PRRs resulted in a significant reduction or tendency of reduction in IL-10 levels released by the moDCs (Fig. 6).

**Figure 6.**
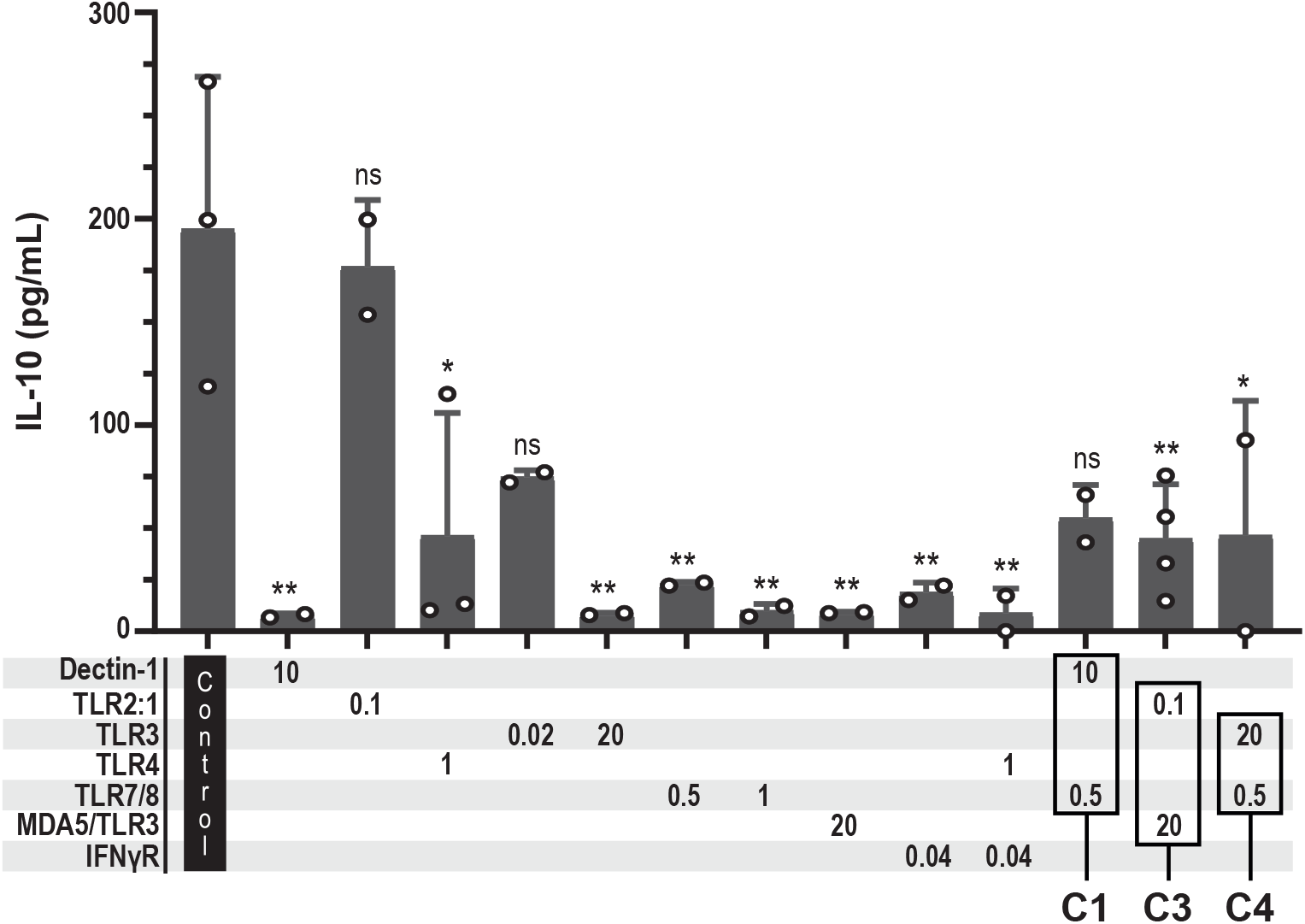
Stimulation of PRRs typically reduced the level of released IL-10 from moDCs. The moDCs were left unstimulated (control) or stimulated for 24 h before IL-10 was quantified in the cell culture supernatants. Each circle presents the concentration from one experiment, and bars represent the mean with SD (n=2-4). The combinations C1, C3, and C4 were tested because they elicited high levels of IL-12p70. The numbers below the bars indicate the concentrations (μg/mL) of the PRR ligands. IL-10 concentrations in media from stimulated moDCs were compared to the concentration in media from unstimulated moDC (control), and statistical significance was evaluated by ANOVA using the Šidák test to correct for multiple comparisons. ns, not significant, *p<0.05, ** p<0.01.

### Stimulation with the TLR2:1 agonist Pam_3_CSK_4_ in combination with transfected poly(I:C) significantly increased the surface expression of CCR7, CD80, CD86, and HLA-DR

The tested ligands and respective PRRs could be divided into three groups depending on whether they triggered production of both IL-12p70 and IFNβ, only one of the two or neither (Table 2). Among these, stimulation of TLR2:1+TLR3 (C2), TLR2:1+MDA5/TLR3 (C3), TLR3+TLR7/8 (C4), or TLR3+NOD2 (C5) resulted in particularly high IL-12p70 and IFNβ production (Table 2). The combination dectin-1+TLR7/8 (C1) stood out as it induced high levels of IL-12p70 but minor-to-non-detectable levels of IFNβ (Table 2). We examined the ability of C1, C3, and C4 to up-regulate the surface expression of four DC activation markers, *i.e*. the chemokine receptor CCR7, the co-stimulatory molecules CD80 and CD86, and the MHC class II molecule HLA-DR (Fig. 7). The expression of the molecules was analysed by flow cytometry in a gated single cell, live population of moDCs (Fig. 7a-d). C3 (TLR2:1+MDA5/TLR3) induced a significant increase in the cell surface expression of all four activation markers (Fig. 7e-i), and C4 (dectin-1+TLR7/8) up-regulated the expression of CD86 (Fig. 7e, h). The remaining test conditions resulted in a tendency to increased surface expression of the activation markers (Fig. 7e-i). Taken together, the combinations C1, C3, and C4 which induced high IL-12p70 production, elicited a trend or significant increase in cell surface expression of the lymph node homing receptor CCR7 and of the molecules CD80, CD86, and HLA-DR which are pivotal in providing signal 1 and signal 2 during T cell activation.

**Figure 7.**
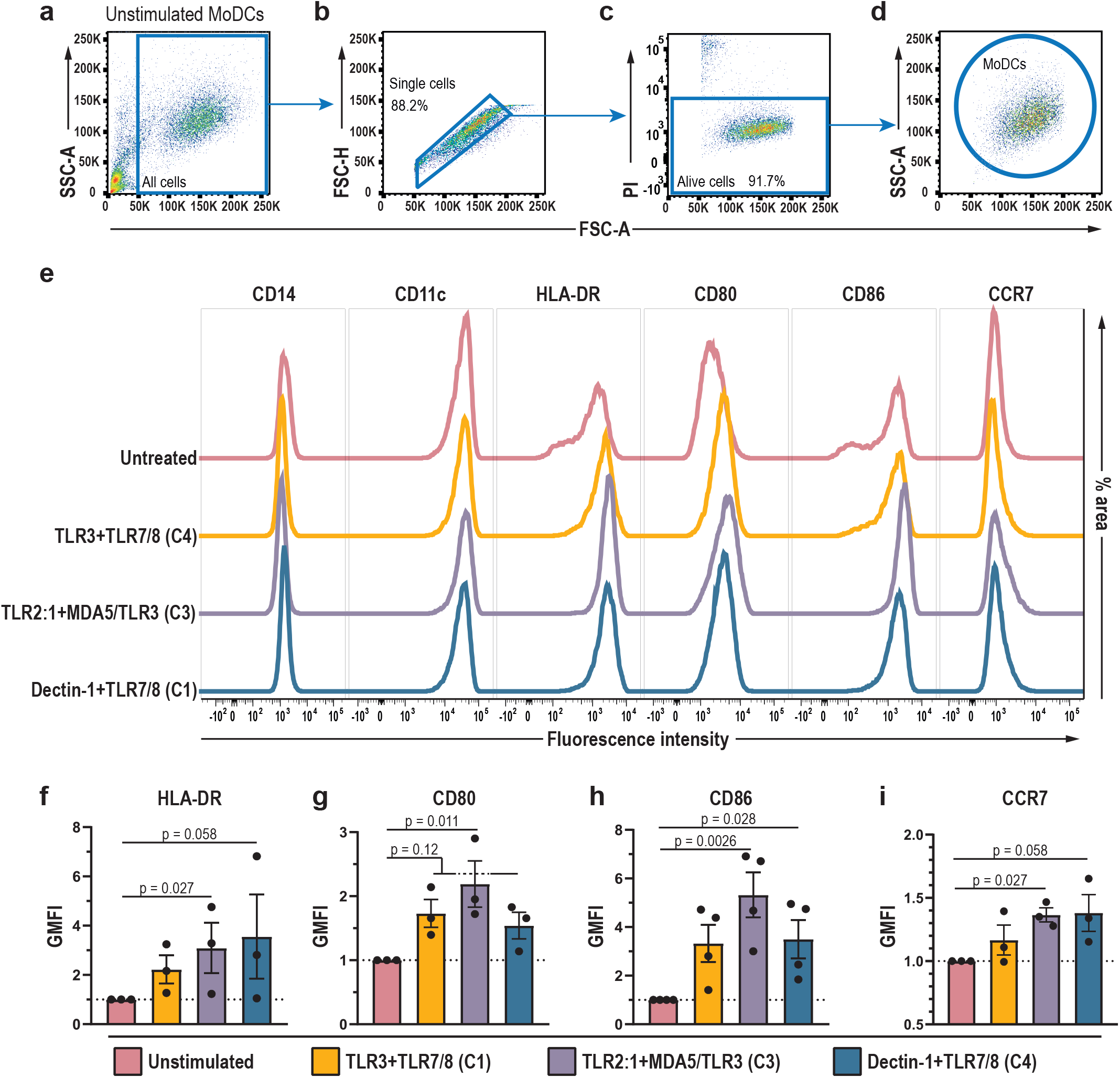
Stimulation of TLR2:1+MDA5/TLR3 increased the surface expression of HLA-DR, CD80, CD86, and CCR7 on moDCs. Cultures of moDCs were stimulated with ligands of TLR3+TLR7/8 (C1), TLR2:1+MDA5/TLR3 (C3), or dectin-1+TLR7/8 (C4) for 24 h before analysis by flow cytometry. Unstimulated moDCs were used as control. **a-d** Before analysis of the surface molecules, single, live moDCs were identified as indicated: **a** Debris was excluded by size (FSC-A), and **b** single cells were selected and doublets excluded in an FSC-A vs. FSC-H plot, before **c**,**d** the live cells, defined as propidium iodide (PI)-negative were selected for analysis of cell surface molecules. **e** Surface expression by moDCs of CD14, CD11c, HLA-DR, CD80, CD86, and CCR7 in a representative sample. CD14 and CD11c showed similar expression levels at all tested conditions whereas PRR stimulation resulted in a significant increase or tendency towards increase in surface expression of the indicated molecules. Geometric mean fluorescence intensity (GMFI) was calculated for **f** HLA-DR, **g** CD80, **h** CD86, and **i** CCR7 and compared to unstimulated moDCs which GMFI was set to 1. The bars indicate mean values with SD, and each dot represents the result from one experiment (n=3-4). P values were calculated by use of an uncorrected Dunn’s Test, and p<0.05 was considered statistically significant.

**Table 2.**
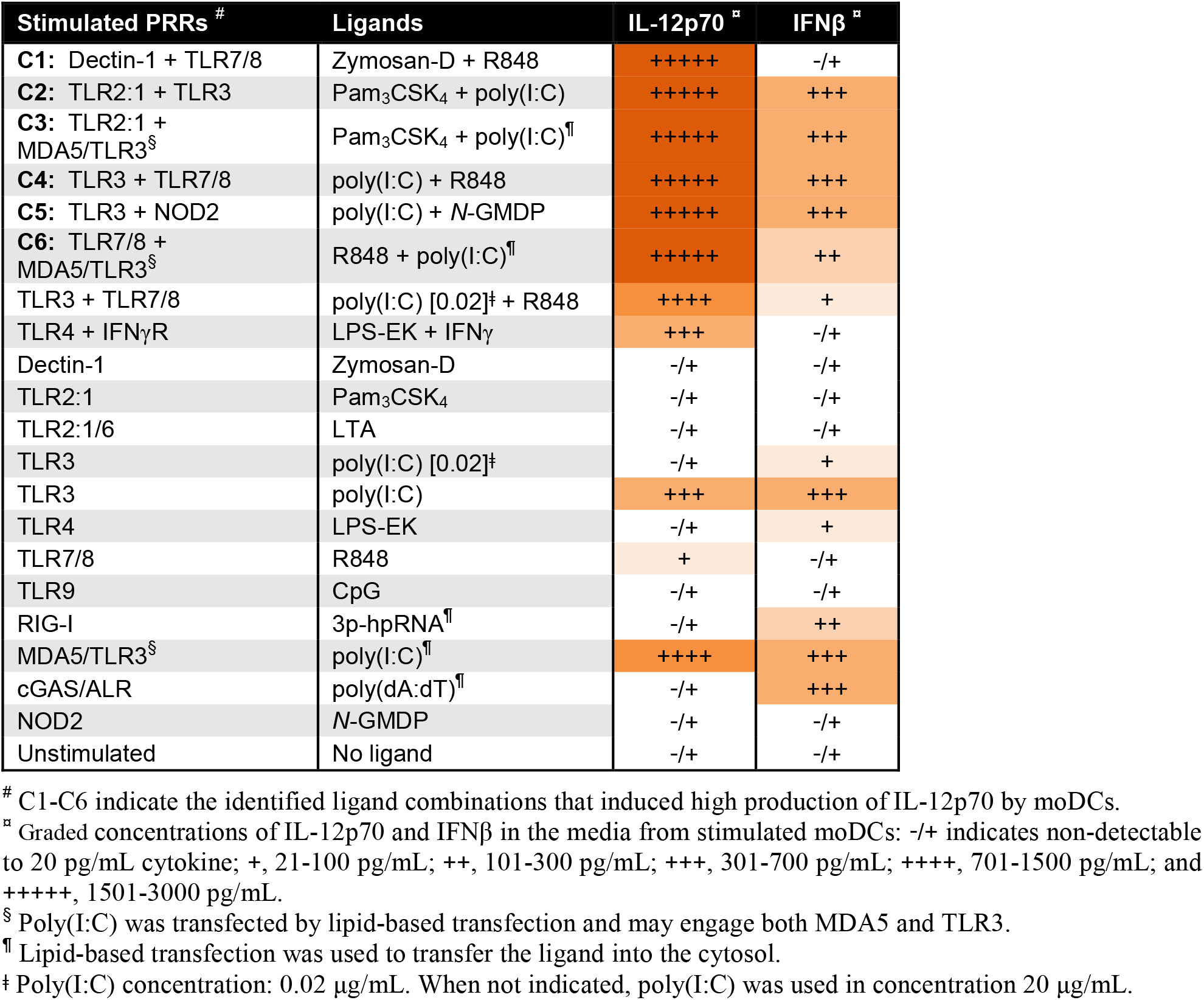
Data summary for the production of IL-12p70 and IFNβ by DCs stimulated with PRR agonists.

## Discussion

In this study, we have stimulated human moDCs with ligands of receptors from all five families of PRRs, with the aim to identify the different PRRs’ ability to induce production of the Th1-polarizing cytokine IL-12p70 and the antiviral cytokine IFNβ. Many of the intracellular PRRs elicited IFNβ production (TLR3, RIG-I, MDA5, and cGAS; but not TLR4, TLR7/8, TLR9, or NOD2), whereas only poly(I:C), a ligand of both TLR3 and MDA5, was able to induce substantial production of IL-12p70. The tested cell surface PRRs dectin-1, TLR2:1, and TLR2:1/6 did neither elicit IL-12p70 nor IFNβ production. All ten tested PRR ligands demonstrated biological activity, and those, that were unable to induce cytokine production as single ligands, showed biological activity in combinations with other PAMPs. Thus, the PRR ligands that we tested could be divided into three groups depending on whether they induced production of both IL-12p70 and IFNβ, only one of the two or neither. Among 26 combinations of different PAMPs only eight combinations, which either contained poly(I:C) or R848 that engage RNA-sensing PRRs, elicited IL-12p70 production. Our data show that IL-12p70 production is subject to strict control in moDCs, and we identified six combinations of PRR ligands that induce high IL-12p70 production and thereby may represent strong adjuvant candidates for vaccines.

### Is stimulation of one PRR sufficient to trigger notable IL-12p70 production – or are two PRRs required?

To stimulate the cytosolic PRRs MDA5 and RIG-I, we transfected DCs with high molecular weight poly(I:C) (1.5-8 kb long) and 5’triphosphate hairpin RNA (89 bp), respectively. Upon stimulation, MDA5 and RIG-I trigger similar signaling pathways, interact with mitochondrial antiviral-signaling protein (MAVS) and activate IRF3, IRF7, and NF-κB^30,39^. However, only poly(I:C), the ligand of MDA5, and not the hairpin 89 bp RNA elicited IL-12p70 production in our experiments. Although it cannot be excluded that MDA5 and RIG-I induce some differences in signaling which potentially could explain the discrepancy in IL-12p70 production, our results made us wonder whether transfected poly(I:C) not only stimulated MDA5 but also TLR3: Some poly(I:C) in the transfection mixture might have been available for stimulation of endosomal TLR3; and *vice versa* when aiming to stimulate TLR3 in the absence of transfection reagent, some poly(I:C) may possibly enter the cytosol and activate MDA5. Thus, stimulation with poly(I:C) (with or without transfection) may engage both TLR3 and MDA5, which potentially could synergize to elicit production of IL-12p70. R848, which induced low levels of IL-12p70 in some experiments, is also an agonist of two receptors, *i.e*. TLR7 and TLR8. Thus, although we could induce IL-12p70 production by a single ligand, poly(I:C) or R848, it remains to be elucidated whether two signals, *i.e*. stimulation of two PRRs were required for the production.

### Combinations of two PAMPs synergistically increase the IL-12p70 level

Here, we found that various combinations of two PAMPs increased the level of IL-12p70 in a synergistic manner. This was different from what we observed for IFNβ, which was produced at similar levels upon stimulation with single ligands and combinations of two ligands. The combination zymosan-D+R848 (C1, dectin-1+TLR7/8) induced the most pronounced effect as it increased the IL-12p70 level 25 to 47-fold compared to the respective single ligands. The combinations C3 (TLR2:1+MDA5/TLR3) and C4 (TLR3+TLR7/8) increased the IL-12p70 level more than 2.5-fold compared to poly(I:C) alone. Furthermore, although C2 (TLR2:1+TLR3), C5 (TLR3+NOD2), or C6 (TLR7/8+MDA5/TLR3) were not compared side by side to their respective single ligands, the combinations appear to have a synergistic effect on the IL-12p70 level. Taken together, our data show that to obtain consistent high levels of IL-12p70 production by moDCs, two signals, *i.e*. stimulation with two different PAMPs are needed. But importantly, any combination of PAMPs does not trigger IL-12p70 production. Among the 26 different combinations we tested, only the combination zymosan-D+R848 (C1, dectin-1+TLR7/8) and the combinations which contained poly(I:C) (except from poly(I:C)+CpG) elicited IL-12p70 production. Thus, all combinations which induced high IL-12p70 production in our experiments contained an agonist of RNA-sensing PRRs (TLR3, MDA5, or TLR7/8).

One of the combinations we tested (0.02 μg/mL poly(I:C)+1 μg/mL R848) has been utilized in clinical cancer vaccine trials^36,40^. Here, we found that modification of the ligand concentrations to 20 μg/mL poly(I:C)+0.05 μg/mL R848 (C4) could increase the level of IL-12p70 >2.5-fold. However, we consider C1, C2, and C3 to be the most promising combinations for cancer immunotherapy. C1 (dectin-1+TLR7/8) stood out as it elicited IL-12p70 but not IFNβ production. C3 (MDA5/TLR3+TLR2:1) induced the highest level of IL-12p70 production and in addition, significantly increased the surface expression of CD80, CD86, HLA-DR, and CCR7.

### Are vita-PAMPs pivotal for IL-12p70 production in the absence of IFNγ?

In accordance with our results, it has been reported that LPS+IFNγ induce IL-12p70 production in DCs, whereas LPS or IFNγ alone does not^41^. Although it is established that PAMPs can trigger IL-12p70 production, the rules governing high production of IL-12p70 remain to be identified. In 2011, the term viability-associated (*vita*)-PAMP was coined to describe PAMPs presenting a signature of microbial life and with the ability to induce immune responses distinct from dead bacteria^42-44^. So far two *vita*-PAMPs have been described, *i.e*. prokaryotic mRNA and cyclic-di-adenosine-monophosphate (c-di-AMP) recognized by TLR8 and STING, respectively^42^. In that context, it is interesting to note that among the ligands we tested, only poly(I:C) and R848 were able to elicit IL-12p70 production. R848 is a ligand of TLR7 and TLR8, and poly(I:C) consisting of dsRNA may mimic a viral infection. Thus, it is tempting to speculate that induction of IL-12p70 production in the absence of IFNγ requires recognition of a PAMP mimicking a live, possibly replicating pathogen. Then, the combination with a second signal, from a PAMP, ensures high production of IL-12p70 and eradication of the live pathogen by type 1 immunity. We propose that exploiting the synergistic effect of combined activation of PRRs for high production of IL-12p70 by DCs may have a great potential for vaccines against pathogens and cancer.

## Abbreviations

ALR: absent in melanoma 2 (AIM2)-like receptor
APC: antigen-presenting cell
CLR: C-type lectin receptor
DC: dendritic cell
ds: double stranded
IFN: interferon
IFNγR: interferon γ receptor
IRF: interferon regulatory factor
LPS: lipopolysaccharide
MDA5: melanoma differentiation-associated protein 5
MDP: muramyl dipeptide
MHC: major histocompatibility complex
moDCs: monocyte-derived dendritic cells
NF-κB: nuclear factor-κB
NLR: nucleotide-binding oligomerization domain (NOD)-like receptor
NOD: nucleotide-binding oligomerization domain
PAMP: pathogen-associated molecular pattern
PRR: pattern recognition receptor
RIG-I: retinoic acid-inducible gene-I
RLR: retinoic acid-inducible gene-I (RIG-I)-like receptor
ss: single stranded
Th1: T helper 1
TLR: toll-like receptor
TRIF: TIR domain-containing adapter molecule 1
*vita*-PAMP: viability-associated PAMP
Zymosan-D: zymosan depleted

## ACKNOWLEDGEMENTS

This work was supported by the South-Eastern Norway Regional Health Authority (grant no. 2018046, and 2022099), the Norwegian Cancer Society (grant no. 198040), and The Research Council of Norway through its Centres of Excellence scheme, project number 262613

## AUTHOR CONTRIBUTIONS

BG and IØ performed experiments. BG, AC, and IØ analyzed and discussed the results, and BG and AC contributed in writing of the manuscript. IØ supervised the study, and wrote the manuscript.

## Supplementary Material

**Supplementary Table 1.**
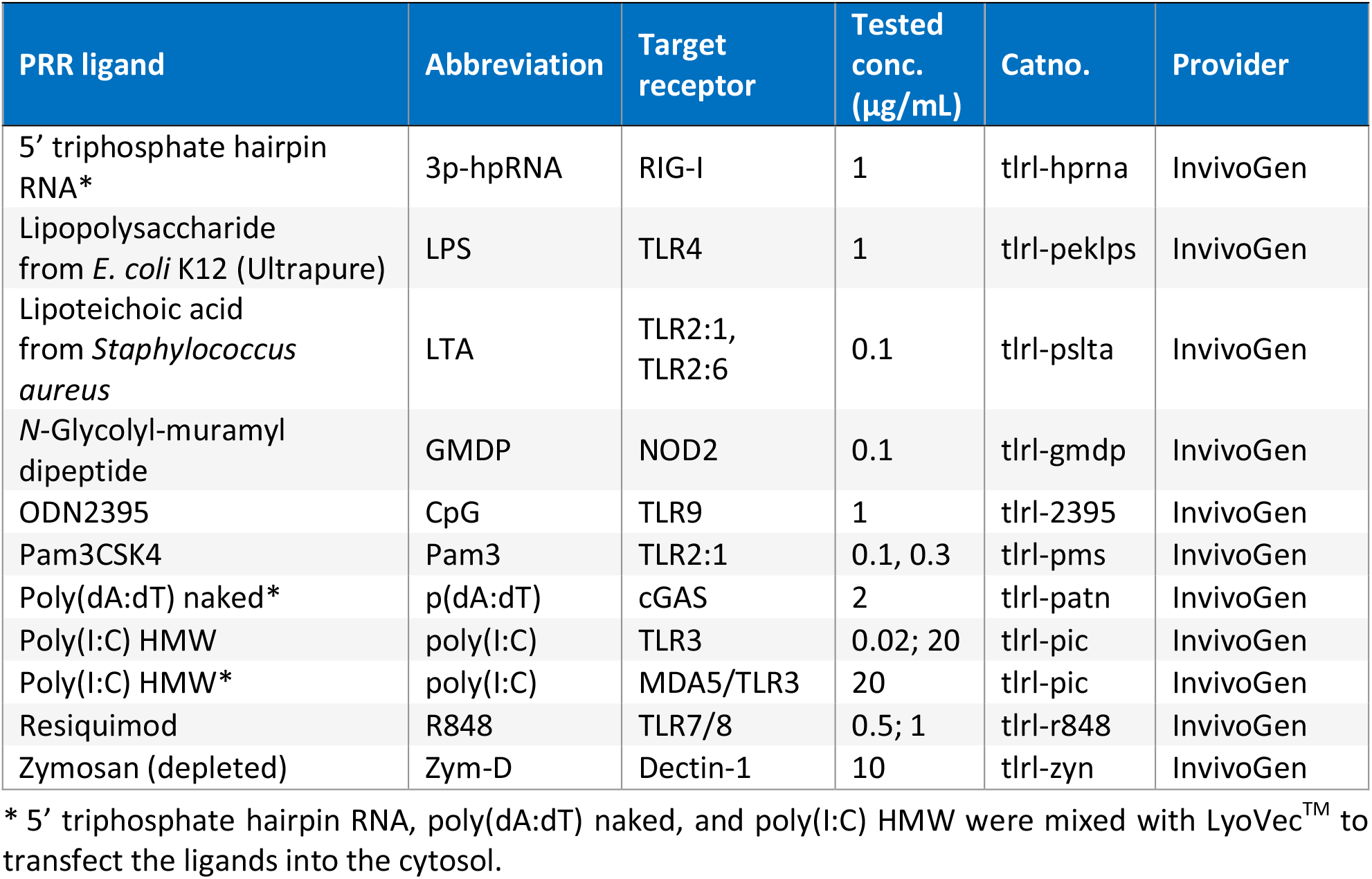
Ligands used to stimulate pattern recognition receptors (PRRs), and ligand abbreviations.

**Supplementary Table 2.** ELISA kits used for quantification of cytokines.

| Item | Provider | Catno. |
| --- | --- | --- |
| DuoSet® Human IL-12 p70 | Bio-Techne | DY1270 |
| DuoSet® Human IFN-β | Bio-Techne | DY814-05 |
| DuoSet® Human IL-10 | Bio-Techne | DY217B |

**Supplementary Table 3.** Primary antibodies used for analysis of expression of cell surface molecules on monocytes and monocyte-derived DCs.

| Specificity | Fluorophore | Clone | Isotype | Final conc. µg/mL | Provider | Catno. |
| --- | --- | --- | --- | --- | --- | --- |
| **CD14** | APC/Cy7 | HCD14 | Mouse IgG1,κ | 20 | BioLegend | 325620 |
| **CD11c** | Pacific Blue | 3.9 | Mouse IgG1,κ | 20 | BioLegend | 301626 |
| **HLA-DR** | BV605 | L243 | Mouse IgG2a,κ | 5 | BioLegend | 307640 |
| **CD80** | PE/Cy7 | 2D10 | Mouse IgG1,κ | 10 | BioLegend | 305217 |
| **CD86** | APC | IT2.2 | Mouse IgG2b,κ | 10 | BioLegend | 305411 |
| **CCR7** | PE/Cy7 | G043H7 | Mouse IgG2a,κ | 10 | BioLegend | 353225 |

**Supplementary Table 4.** Isotype- and fluorophore-matched, irrelevantly targeted antibodies used as negative controls.

| Fluorophore | Isotype | Clone | Specificity | Provider | Catno. |
| --- | --- | --- | --- | --- | --- |
| **APC/Cy7** | Mouse IgG1,$\kappa$ | MOPC-21 | Unknown | BioLegend | 400128 |
| **Pacific Blue** | Mouse IgG1,$\kappa$ | MOPC-21 | Unknown | BioLegend | 400151 |
| **BV605** | Mouse IgG2a,$\kappa$ | MOPC-173 | Unknown | BioLegend | 400270 |
| **PE/Cy7** | Mouse IgG1,$\kappa$ | MOPC-21 | Unknown | BioLegend | 400126 |
| **APC** | Mouse IgG2b,$\kappa$ | MG2b-57 | Trinitrophenol + KLH | BioLegend | 401210 |
| **PE/Cy7** | Mouse IgG2a,$\kappa$ | MOPC-173 | Unknown | BioLegend | 400231 |

**Supplementary Table 5.** Increased concentrations of the agonists of dectin-1, TLR2:1, or TLR2:6 did not trigger IL-12p70 production.

| Stimulated PRR | Ligand | Ligand concentration (pg/mL) | Absorbance |  | IL-12p70 (pg/mL) <sup>a</sup> |
| --- | --- | --- | --- | --- | --- |
| **Unstimulated** | None | 0 | 0.081 | 0.078 | ND |
| **Dectin-1** | Zym-D | 1 | 0.083 | 0.081 | ND |
|  |  | 10 | 0.083 | 0.081 | ND |
|  |  | 20 | 0.080 | 0.083 | ND |
| **TLR2:1** | Pam<sub>3</sub>CSK<sub>4</sub> | 0.5 | 0.088 | 0.082 | ND |
|  |  | 1 | 0.080 | 0.080 | ND |
|  |  | 10 | 0.083 | 0.078 | ND |
| **TLR2:1; TLR2:6** | LTA | 0.5 | 0.083 | 0.078 | ND |
|  |  | 1 | 0.079 | 0.079 | ND |
|  |  | 10 | 0.084 | 0.080 | ND |
|  |  | 100 | 0.082 | 0.081 | ND |
| **MDA5/TLR3** | poly(I:C)* | 20 | 1.353 | 1.455 | 1470.5 |

ND, not detectable
\* Poly(I:C) was transfected into the cytosol by LyoVec<sup>TM</sup>, a lipid-based transfection reagent. The ligand was included as positive control.
<sup>a</sup> Amounts of IL-12p70 (pg/mL) in supernatants from monocyte-derived DCs stimulated with the indicated ligand concentrations.

**Supplementary Table 6.** Increased concentrations of the ligands of dectin-1, TLR2:1, or TLR2:6 did not trigger IFNβ production.

| Stimulated PRR | Ligand | Ligand concentration (pg/mL) | IFNβ (pg/mL) <sup>‡</sup> | |
| --- | --- | --- | --- | --- |
| **Unstimulated** | None | 0 | 7.1 | 7.5 |
| **Dectin-1** | Zym-D | 1 | 7.1 | 7.4 |
|  |  | 10 | 6.9 | 7.0 |
|  |  | 20 | 6.9 | 6.7 |
| **TLR2:1** | Pam<sub>3</sub>CSK<sub>4</sub> | 0.5 | 7.1 | 7.6 |
|  |  | 1 | 6.9 | 7.1 |
|  |  | 10 | 7.0 | 7.2 |
| **TLR2:1, TLR2:6** | LTA | 0.5 | 7.1 | 7.0 |
|  |  | 1 | 7.1 | 7.1 |
|  |  | 10 | 7.1 | 7.8 |
|  |  | 100 | 7.1 | 7.2 |
| **MDA5/TLR3** | poly(I:C) * | 20 | 261.9 | 289.2 |

\* Poly(I:C) was transfected into the cytosol by LyoVec<sup>™</sup>, a lipid-based transfection reagent. The ligand was included as positive control.
<sup>‡</sup> Amounts of IFNβ (pg/mL) in supernatants from monocyte-derived DCs stimulated with the indicated ligand concentrations.

**Supplementary Figure 1.**
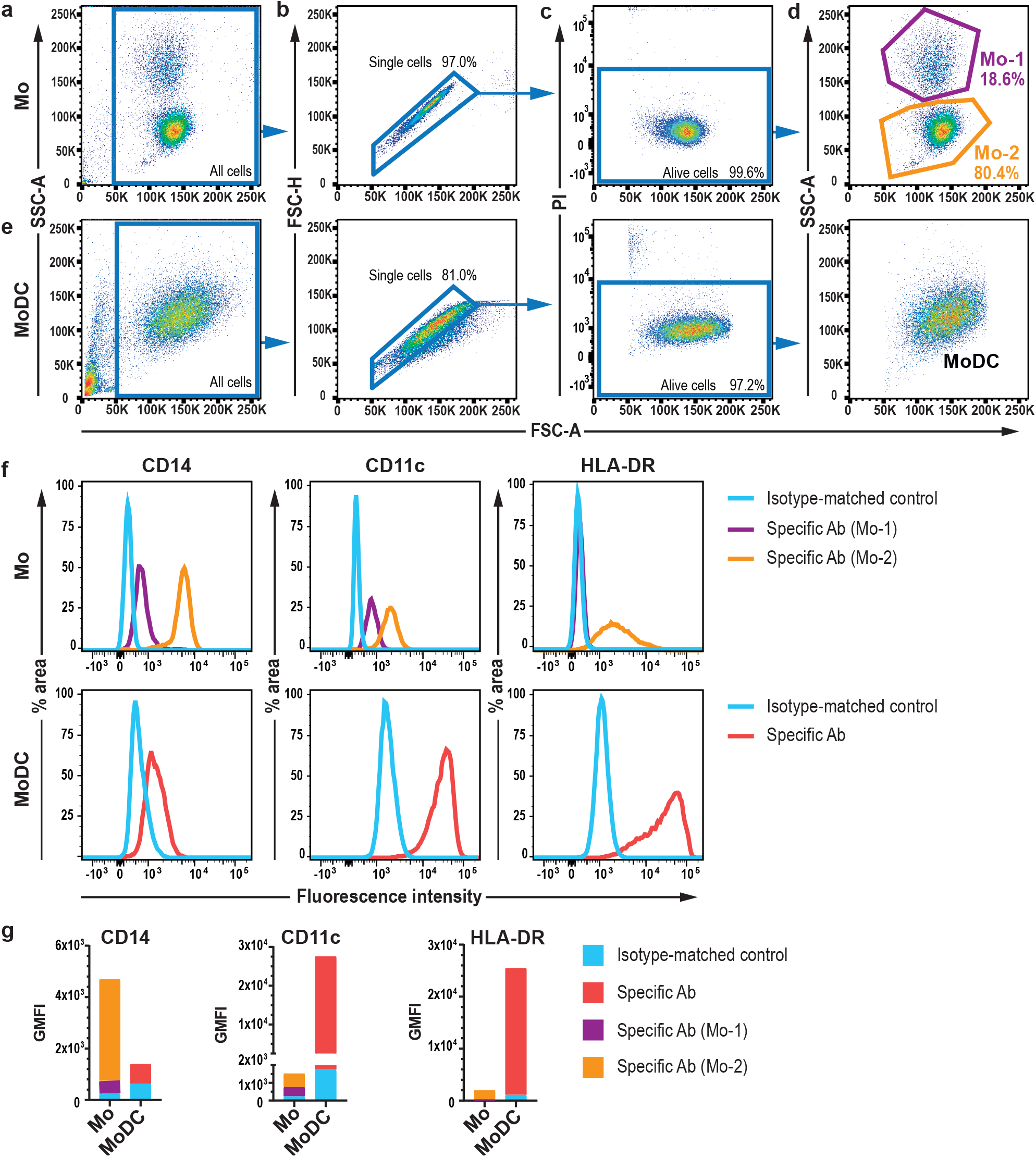
Gating and flow cytometry analysis of monocytes and monocyte-derived DCs. Positively selected, freshly isolated CD14+ monocytes and monocytes differentiated for 5 days by GM-CSF and IL-4 into monocyte-derived dendritic cells (moDC) were gated and analysed by flow cytometry as indicated: **a** The monocytes (Mo) were first gated based on size and complexity in a forward scatter (FSC-A) and side scatter (SSC-A) plot; **b** Next, a FSC-A and FSC-H plot was used to gate single cells and exclude doublets; **c** Dead cells were removed by their uptake of propidium iodide (PI); **d** Gated live, single cells distributed in two populations, Mo-1 and Mo-2, which constituted 18.6 and 80.4 % respectively, of the gated monocytes. **e** The same gating strategy was used for moDCs, and gated live, single moDCs distributed in one population in the FSC-A and SSC-A plot. **f** Mo-1 showed low CD14 and CD11c expression, and was negative for HLA-DR. Mo-2 on the other hand, was positive for CD14, CD11c, and HLA-DR. The moDCs showed reduced expression of CD14 compared to Mo-2, and increased expression of CD11c and HLA-DR compared to both Mo-1 and Mo-2. **g** The graphs show geometric mean fluorescence intensity (GMFI) for CD14, CD11c, and HLA-DR calculated from the experiment shown in f. The specific antibodies that were used: anti-CD14, clone HCD14; anti-CD11c, clone 3.9; and anti-HLA-DR, clone L243. The antibodies were used alongside isotype- and concentration-matched controls.

**Supplementary Figure 2.**
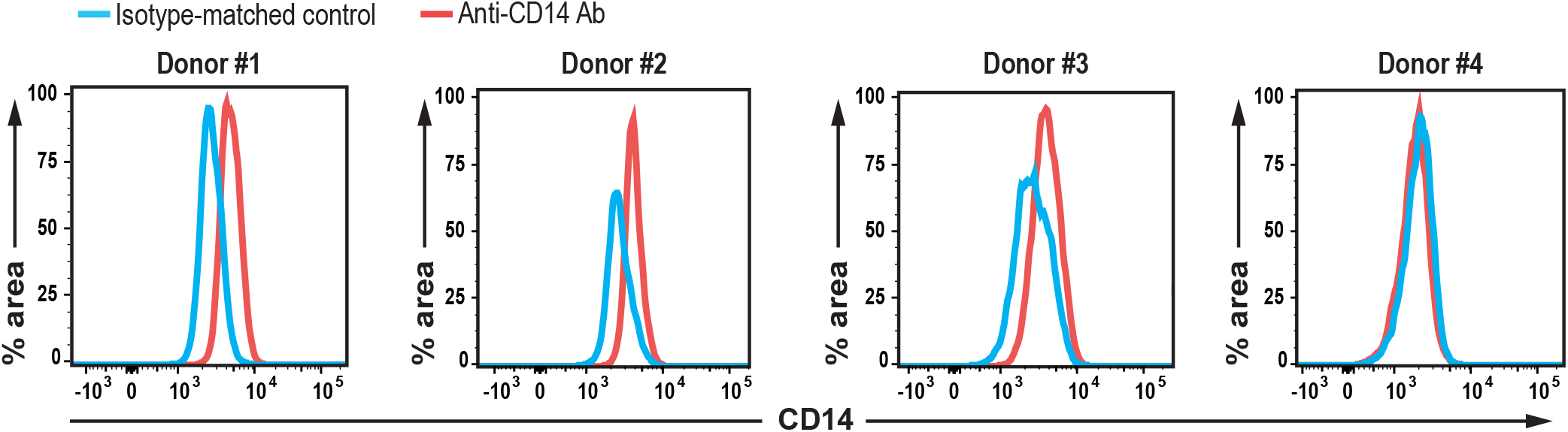
All included blood donors gave rise to monocyte-derived DCs with low CD14 expression. Monocyte-derived dendritic cells (moDCs) were differentiated with GM-CSF (100 ng/mL) and IL-4 (20 ng/mL) for 5 days before flow cytometry analysis of CD14. The moDCs were gated as indicated in Supplementary Figure 1 before evaluation of CD14 expression. The control antibody (blue line, clone MOPC-21) matched the isotype and concentration of the anti-CD14 antibody (red line, clone HCD14). MoDCs generated from different donors (Donor 1-4), showed low to no expression of CD14, suggesting that the monocytes were successfully differentiated into moDCs. The proportion of CD14+ moDCs varied from 13.8-31.8 % with an average value of 22 % across the donors.

## References

1. Burnet F. The clonal selection theory of acquired immunity.: Nashville, TN: Vanderbilt University Press.; 1959.

2. Zinkernagel RM. Restriction by H-2 gene complex of transfer of cell-mediated immunity to Listeria monocytogenes. Nature 1974;251:230–3.

3. Cunningham AJ, Lafferty KJ. A simple conservative explanation of the H-2 restriction of interactions between lymphocytes. Scand J Immunol 1977;6:1–6.

4. Linsley PS, Clark EA, Ledbetter JA. T-cell antigen CD28 mediates adhesion with B cells by interacting with activation antigen B7/BB-1. Proceedings of the National Academy of Sciences of the United States of America 1990;87:5031–5.

5. Corthay A. A three-cell model for activation of naive T helper cells. Scand J Immunol 2006;64:93–6.

6. Hsieh CS, Macatonia SE, Tripp CS, Wolf SF, O’Garra A, Murphy KM. Development of TH1 CD4+ T cells through IL-12 produced by Listeria-induced macrophages. Science 1993;260:547–9.

7. Curtsinger JM, Schmidt CS, Mondino A, et al. Inflammatory cytokines provide a third signal for activation of naive CD4+ and CD8+ T cells. Journal of immunology 1999;162:3256–62.

8. Scott P. IL-12: initiation cytokine for cell-mediated immunity. Science 1993;260:496–7.

9. Ossendorp F, Mengede E, Camps M, Filius R, Melief CJ. Specific T helper cell requirement for optimal induction of cytotoxic T lymphocytes against major histocompatibility complex class II negative tumors. The Journal of experimental medicine 1998;187:693–702.

10. Corthay A, Skovseth DK, Lundin KU, et al. Primary antitumor immune response mediated by CD4+ T cells. Immunity 2005;22:371–83.

11. Qin Z, Blankenstein T. CD4+ T cell--mediated tumor rejection involves inhibition of angiogenesis that is dependent on IFN gamma receptor expression by nonhematopoietic cells. Immunity 2000;12:677–86.

12. Pulendran B, p SA, O’Hagan DT. Emerging concepts in the science of vaccine adjuvants. Nat Rev Drug Discov 2021.

13. Napolitani G, Rinaldi A, Bertoni F, Sallusto F, Lanzavecchia A. Selected Toll-like receptor agonist combinations synergistically trigger a T helper type 1-polarizing program in dendritic cells. Nat Immunol 2005;6:769–76.

14. Ma X, Chow JM, Gri G, et al. The interleukin 12 p40 gene promoter is primed by interferon gamma in monocytic cells. The Journal of experimental medicine 1996;183:147–57.

15. Goriely S, Molle C, Nguyen M, et al. Interferon regulatory factor 3 is involved in Toll-like receptor 4 (TLR4)- and TLR3-induced IL-12p35 gene activation. Blood 2006;107:1078–84.

16. Laderach D, Compagno D, Danos O, Vainchenker W, Galy A. RNA interference shows critical requirement for NF-kappa B p50 in the production of IL-12 by human dendritic cells. Journal of immunology 2003;171:1750–7.

17. Zhu Q, Egelston C, Vivekanandhan A, et al. Toll-like receptor ligands synergize through distinct dendritic cell pathways to induce T cell responses: implications for vaccines. Proceedings of the National Academy of Sciences of the United States of America 2008;105:16260–5.

18. Kobayashi M, Fitz L, Ryan M, et al. Identification and purification of natural killer cell stimulatory factor (NKSF), a cytokine with multiple biologic effects on human lymphocytes. The Journal of experimental medicine 1989;170:827–45.

19. Rogge L, Papi A, Presky DH, et al. Antibodies to the IL-12 receptor beta 2 chain mark human Th1 but not Th2 cells in vitro and in vivo. Journal of immunology 1999;162:3926–32.

20. Wolf SF, Temple PA, Kobayashi M, et al. Cloning of cDNA for natural killer cell stimulatory factor, a heterodimeric cytokine with multiple biologic effects on T and natural killer cells. Journal of immunology 1991;146:3074–81.

21. Snijders A, Hilkens CM, van der Pouw Kraan TC, Engel M, Aarden LA, Kapsenberg ML. Regulation of bioactive IL-12 production in lipopolysaccharide-stimulated human monocytes is determined by the expression of the p35 subunit. Journal of immunology 1996;156:1207–12.

22. Becker C, Wirtz S, Blessing M, et al. Constitutive p40 promoter activation and IL-23 production in the terminal ileum mediated by dendritic cells. J Clin Invest 2003;112:693–706.

23. Janeway CA, Jr. Approaching the asymptote? Evolution and revolution in immunology. Cold Spring Harbor symposia on quantitative biology 1989;54 Pt 1:1–13.

24. Fitzgerald KA, Kagan JC. Toll-like Receptors and the Control of Immunity. Cell 2020;180:1044–66.

25. Del Fresno C, Iborra S, Saz-Leal P, Martinez-Lopez M, Sancho D. Flexible Signaling of Myeloid C-Type Lectin Receptors in Immunity and Inflammation. Frontiers in immunology 2018;9:804.

26. Feinberg H, Mitchell DA, Drickamer K, Weis WI. Structural basis for selective recognition of oligosaccharides by DC-SIGN and DC-SIGNR. Science 2001;294:2163–6.

27. Bermejo-Jambrina M, Eder J, Helgers LC, et al. C-Type Lectin Receptors in Antiviral Immunity and Viral Escape. Frontiers in immunology 2018;9:590.

28. Taylor PR, Tsoni SV, Willment JA, et al. Dectin-1 is required for beta-glucan recognition and control of fungal infection. Nat Immunol 2007;8:31–8.

29. Speakman EA, Dambuza IM, Salazar F, Brown GD. T Cell Antifungal Immunity and the Role of C-Type Lectin Receptors. Trends Immunol 2020;41:61–76.

30. Rehwinkel J, Gack MU. RIG-I-like receptors: their regulation and roles in RNA sensing. Nature reviews Immunology 2020;20:537–51.

31. Qureshi N, Takayama K, Ribi E. Purification and structural determination of nontoxic lipid A obtained from the lipopolysaccharide of Salmonella typhimurium. J Biol Chem 1982;257:11808–15.

32. Evans JT, Cluff CW, Johnson DA, Lacy MJ, Persing DH, Baldridge JR. Enhancement of antigen-specific immunity via the TLR4 ligands MPL adjuvant and Ribi.529. Expert Rev Vaccines 2003;2:219–29.

33. Campbell JD. Development of the CpG Adjuvant 1018: A Case Study. Methods Mol Biol 2017;1494:15–27.

34. Garcon N, Chomez P, Van Mechelen M. GlaxoSmithKline Adjuvant Systems in vaccines: concepts, achievements and perspectives. Expert Rev Vaccines 2007;6:723–39.

35. Zobywalski A, Javorovic M, Frankenberger B, et al. Generation of clinical grade dendritic cells with capacity to produce biologically active IL-12p70. J Transl Med 2007;5:18.

36. Subklewe M, Geiger C, Lichtenegger FS, et al. New generation dendritic cell vaccine for immunotherapy of acute myeloid leukemia. Cancer Immunol Immunother 2014;63:1093–103.

37. Le Naour J, Galluzzi L, Zitvogel L, Kroemer G, Vacchelli E. Trial watch: TLR3 agonists in cancer therapy. Oncoimmunology 2020;9:1771143.

38. Sallusto F, Lanzavecchia A. Efficient presentation of soluble antigen by cultured human dendritic cells is maintained by granulocyte/macrophage colony-stimulating factor plus interleukin 4 and downregulated by tumor necrosis factor alpha. The Journal of experimental medicine 1994;179:1109–18.

39. Onomoto K, Onoguchi K, Yoneyama M. Regulation of RIG-I-like receptor-mediated signaling: interaction between host and viral factors. Cell Mol Immunol 2021;18:539–55.

40. Lichtenegger FS, Mueller K, Otte B, et al. CD86 and IL-12p70 are key players for T helper 1 polarization and natural killer cell activation by Toll-like receptor-induced dendritic cells. PLoS One 2012;7:e44266.

41. Abdi K, Singh N, Matzinger P. T-cell control of IL-12p75 production. Scand J Immunol 2006;64:83–92.

42. Sander LE, Davis MJ, Boekschoten MV, et al. Detection of prokaryotic mRNA signifies microbial viability and promotes immunity. Nature 2011;474:385–9.

43. Moretti J, Roy S, Bozec D, et al. STING Senses Microbial Viability to Orchestrate Stress-Mediated Autophagy of the Endoplasmic Reticulum. Cell 2017;171:809–23 e13.

44. Ugolini M, Gerhard J, Burkert S, et al. Recognition of microbial viability via TLR8 drives TFH cell differentiation and vaccine responses. Nat Immunol 2018;19:386–96.

45. Brown GD, Herre J, Williams DL, Willment JA, Marshall AS, Gordon S. Dectin-1 mediates the biological effects of beta-glucans. The Journal of experimental medicine 2003;197:1119–24.

46. Ikeda Y, Adachi Y, Ishii T, Miura N, Tamura H, Ohno N. Dissociation of Toll-like receptor 2-mediated innate immune response to Zymosan by organic solvent-treatment without loss of Dectin-1 reactivity. Biol Pharm Bull 2008;31:13–8.

47. Brown GD, Gordon S. Immune recognition. A new receptor for beta-glucans. Nature 2001;413:36–7.

